# Brain Representations of Affective Valence and Intensity in Sustained Pleasure and Pain

**DOI:** 10.1101/2023.06.08.544230

**Authors:** Soo Ahn Lee, Jae-Joong Lee, Jisoo Han, Myunghwan Choi, Tor D. Wager, Choong-Wan Woo

**Author notes:** Co-corresponding authors Please address correspondence to: Jae-Joong Lee, Choong-Wan Woo, Department of Biomedical Engineering, Sungkyunkwan University Center for Neuroscience Imaging Research, Institute for Basic Science Suwon 16419, Republic of Korea.

## Abstract

Pleasure and pain are two opposites that compete and influence each other, implying the existence of brain systems that integrate them to generate modality-general affective experiences. Here, we examined the brain’s general affective codes (i.e., affective valence and intensity) across sustained pleasure and pain through an fMRI experiment (*n* = 58). We found that the distinct sub-populations of voxels within the ventromedial and lateral prefrontal cortices, the orbitofrontal cortex, the anterior insula, and the amygdala were involved in decoding affective valence versus intensity, which was replicated in an independent test dataset (*n* = 62). The affective valence and intensity models were connected to distinct large-scale brain networks—the intensity model to the ventral attention network and the valence model to the limbic and default mode networks. Overall, this study identified the brain representations of affective valence and intensity across pleasure and pain, promoting the systems-level understanding of human affective experiences.

## INTRODUCTION

Pain and pleasure are two representative emotional entities of negative and positive affect (*1*). They compete and interact with each other to generate the subjective interpretation of current hedonic status (*2*). For instance, pleasant stimuli can reduce pain, while painful stimuli can reduce pleasure. In addition, pain relief is often perceived as pleasant, and the absence of pleasure, also known as anhedonia, is common in patients with chronic pain. This implies that there exist brain systems that integrate pain and pleasure—although painful and pleasant sensations are processed through distinct spinal and peripheral circuits (*3, 4*), they should ultimately be integrated into modality-general affective experiences (*5, 6*) in higher-level core affective brain systems (*7, 8*). This idea has been supported by a large body of literature showing the overlap in the brain regions involved in pain and pleasure (*9*). Many of the overlapping brain regions are part of opioid systems that play important roles in hedonic feelings and motivation (*10*) and the interaction between pain and pleasure (*9, 11*). However, most of the previous studies have been conducted for either pain or pleasure separately, and a direct comparison of their neural representations within the same individuals is lacking. Although some human functional Magnetic Resonance Imaging (fMRI) studies have identified a set of brain regions that respond to both positive and negative emotions (*12–14*), these studies have used non-painful aversive stimuli rather than painful stimuli. A few animal studies have reported that the amygdala (*15*) and the anterior cingulate cortex (*16*) contain neurons that respond to both pain and pleasure, but these findings have been limited to specific local regions, and their generalizability to humans has not been well established.

In addition, it remains elusive how brain regions that are activated by both pain and pleasure differentially encode each type of affect. One possibility is that these brain regions represent *affective valence*, which ranges from negative to positive on a hedonic continuum where pain and pleasure show a marked difference (*17*). However, a growing number of studies have found some neural populations that also respond to both positive and negative valence, which has been referred to as the valence-general affective workspace (*18*), unsigned valence (*19*), arousal (*7, 20*) and salience (*21, 22*). We refer to this as *affective intensity,* which reflects the magnitudes of positive and negative feelings and is proposed as another fundamental dimension of core affect. Both affective valence (referred to here as “valence”) and affective intensity (referred to here as “intensity”) are important for decision-making and survival because they influence an organism’s approach or avoidance responses to a stimulus or situation and its perceived importance (*23*), potentially involving distinct neural populations (*24*). However, the brain representations of these general affective codes, i.e., affective valence and intensity, have yet to be identified at the systems level.

In this study, we examined the brain representations of affective valence and intensity across sustained pleasure and pain fMRI with the information mapping (*25*) and predictive modeling approaches (*26*). More specifically, our key research questions include: (1) Which brain regions contain information about pleasure and pain? Among them, which brain regions overlap? (2) Can we identify predictive models of affective valence and intensity within the overlapping brain regions? (3) Which large-scale brain networks are correlated with these predictive models? To answer these questions, we conducted an fMRI experiment (**Study 1**, *n* = 58). Participants continuously rated their subjective pleasantness and unpleasantness while experiencing sustained pain and pleasure. We induced sustained pain and pleasure by delivering capsaicin and chocolate fluids, respectively, into the participants’ oral cavities via a mouthpiece (Fig. 1A). During the fMRI scans, which lasted 14.5 minutes and included two-times fluid deliveries, participants provided continuous ratings using a modified version of the general Labeled Magnitude Scale (gLMS) (*27*). Water was delivered at all other times (Fig. 1B).

**Fig. 1.**
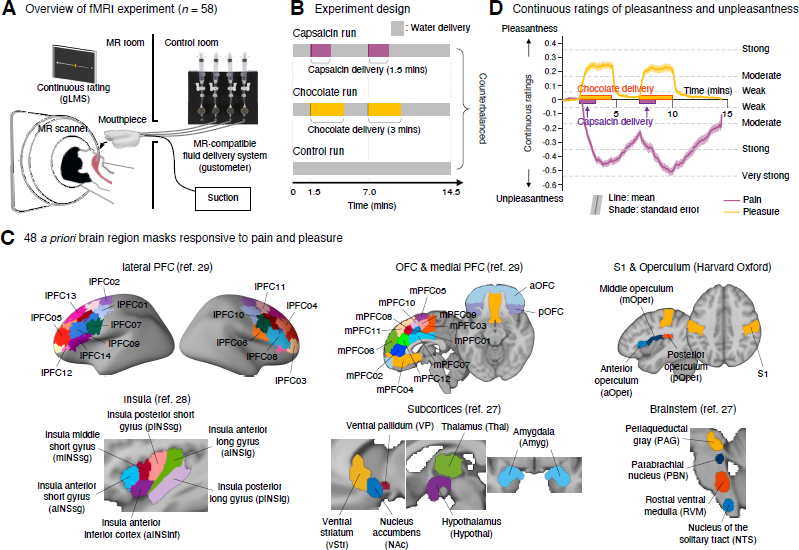
Experimental overview and regions of interest. **(A)** The overview of the fMRI experiment. We delivered fluids using an MR-compatible fluid delivery system—a gustometer. We also used a suction device to remove fluids from a participant’s oral cavity during the experiment. While experiencing pain and pleasure induced by the capsaicin and chocolate fluids, participants were asked to continuously rate their subjective feelings of pleasantness and unpleasantness using the general Labeled Magnitude Scale (gLMS). **(B)** There were three different experimental conditions, including the ‘capsaicin’ (pain), ‘chocolate’ (pleasure), and ‘control’ (neutral) conditions. The capsaicin or chocolate fluid was delivered twice in the middle of the scan. The capsaicin fluid was delivered for a duration of 1.5 mins each, and the chocolate fluid was delivered for a duration of 3 mins each. All runs lasted 14.5 mins, and the order of the conditions was counterbalanced across the participants. **(C)** The locations of 48 *a priori* regions-of-interest (ROIs). The ROIs were selected based on previous literature, as provided in **Fig. S1**. **(D)** The continuous ratings for the capsaicin and chocolate conditions (*n* = 58). The solid lines indicate group average ratings (purple for pain, yellow for pleasure), and the shading indicates standard errors of the mean (s.e.m.).

We first used the multivariate pattern-based information mapping approach to identify the brain regions containing information about dynamic changes in pleasantness-unpleasantness ratings. Among the brain regions which showed significant decoding performances for both pain and pleasure, the overlapping brain regions were the ventromedial and lateral prefrontal cortices, the posterior orbitofrontal cortex, the ventral anterior insula, and the amygdala, which largely correspond to the neural reference space of core affect (*7*). We then developed predictive models of valence and intensity using these 7 overlapping brain regions to identify and compare the fine-grained brain representations of each affective dimension (*18*). Both the valence and intensity models showed significant prediction performance in the training and independent test datasets (**Study 2**, *n* = 62). The important predictive features of these models included largely non-overlapping subregions distributed across the 7 brain regions. The co-localized, but distinct brain representations of valence and intensity were correlated with distinct large-scale brain networks—the valence model was correlated with the limbic and default mode networks, whereas the intensity model was correlated with the ventral attention network.

## RESULTS

### Identifying the brain regions containing information about sustained pleasure and pain

To address the first research question (“Which brain regions contain information about pleasure and pain? Among them, which brain regions overlap?”), we identified a set of brain regions whose activation patterns are predictive of pleasantness-unpleasantness ratings during the ‘chocolate’ (i.e., pleasure) or ‘capsaicin’ (i.e., pain) condition. To minimize false positives, we limited our search to the 48 *a priori* regions-of-interest (ROIs) known to respond to either pain or pleasure in the previous literature (Fig. 1C). The ROIs include brainstem and midbrain regions including the nucleus of the solitary tract, rostral ventral medulla, parabrachial nucleus, and periaqueductal gray (*28*), subcortical regions including the amygdala, ventral pallium, ventral striatum, nucleus accumbens, thalamus, and hypothalamus (*28*), insular cortex regions including insula anterior inferior cortex, insula anterior/middle/posterior short gyri, and insula anterior/posterior long gyri (*29*), primary somatosensory cortex, anterior/middle/posterior operculum, anterior/posterior orbitofrontal cortices (OFCs), 12 parcellations of the medial prefrontal cortex (mPFC01∼12) (*30*), and 14 parcellations of the lateral prefrontal cortex (lPFC01∼14) (*30*). **Fig. S1** provides the list of previous studies that reported these ROIs’ responses to pain or pleasure stimuli.

For each of these ROIs, we trained fMRI pattern-based decoders to predict pleasantness-unpleasantness ratings for the pain or pleasure condition separately. Fig. 1D shows the overall patterns of pleasantness-unpleasantness ratings for the capsaicin and chocolate conditions, which served as the outcome variable in this pattern-based information mapping. The rating patterns clearly show two peaks responding to two deliveries of capsaicin and chocolate, which is an important design feature to improve the specificity of predictive models. We provided the details of the predictive modeling in **Methods** and **Fig. S2**, but briefly here, we used beta estimates of 34-time bins from a general linear model (GLM) of fMRI data (i.e., 25 seconds per time-bin) to enhance the signal-to-noise ratio and remove nuisance effects. We applied principal component regression (PCR) on the ‘capsaicin’ and ‘control’ condition data to develop pain models and on the ‘chocolate’ and ‘control’ condition data to develop pleasure models. To account for the different numbers of voxels in each ROI, we used the same number of principal components (13 PCs) across ROIs (for details of how we selected the number of PC, see **Methods**). We calculated model performance with the averaged within-individual correlation between predicted and actual ratings using leave-one-subject-out cross-validation (LOSO-CV).

A total of 7 among the 48 ROIs showed significant decoding performance for both pain and pleasure (Fig. 2A), which include the amygdala, the insula anterior inferior cortex (which is also called ventral anterior insula; below, we added alternative region names in the parenthesis), mPFC04 (ventromedial PFC), the posterior OFC, lPFC05, lPFC09, and lPFC14 (ventro-to dorso-lateral PFC). Their mean within-individual correlations between the actual and predicted ratings were *r* = 0.11-0.18, mean within-individual mean squared error (mse) = 0.052-0.054, *P* = 0.00001-0.0045 for predicting pain, and mean *r* = 0.09-0.13, mean mse = 0.021-0.022, *P* = 0.0005-0.0105 for predicting pleasure. These were significant after the correction for multiple comparisons with False Discovery Rate (FDR) *q* < 0.05, bootstrap test, two-tailed. Many of these regions are among the brain structures that have been implicated in the common neurobiology of pain and pleasure (*9*) and substantially overlap with the hypothetical neural reference space of core affect (i.e., ‘affective workspace’) (*7*). The brain regions important only for the prediction of pain (i.e., pain-unique) included the anterior OFC, mPFC07 (mid-cingulate cortex), lPFC03 (frontal pole), and lPFC13 (pre-supplementary motor area). The brain regions important only for the prediction of pleasure (i.e., pleasure-unique) included the insula anterior and posterior long gyri, mPFC01 (a dorsal part of the mid-cingulate cortex), and lPFC11 (supplementary motor area).

**Fig. 2.**
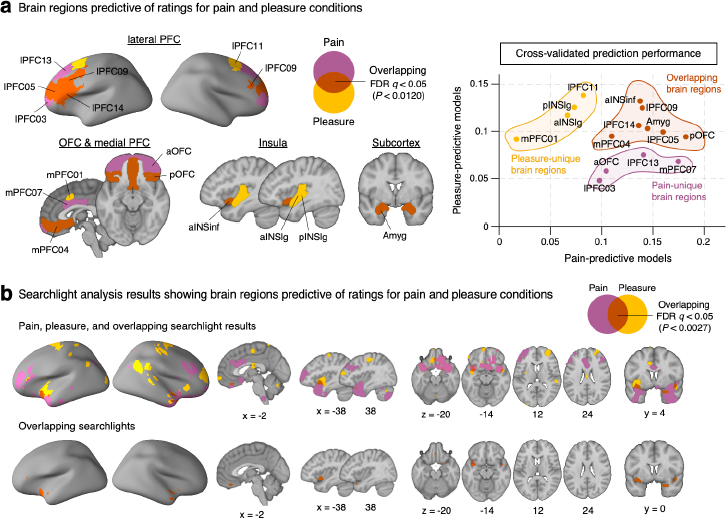
Information mapping results. **(A)** Left: Brain regions that showed significant prediction performances (FDR *q* < 0.05) at predicting the pleasantness-unpleasantness ratings for the pain and pleasure conditions. The overlapping brain regions are colored in reddish brown. Right: A scatter plot showing the cross-validated prediction performances. aINSinf, insula anterior inferior cortex; aINSlg, insula anterior long gyrus; pINSlg, insula posterior long gyrus; Amyg, amygdala; aOFC, anterior orbitofrontal cortex; pOFC, posterior orbitofrontal cortex; lPFC, lateral prefrontal cortex; mPFC, medial prefrontal cortex. **(B)** Top: Searchlight voxels that showed significant prediction performances (FDR *q* < 0.05). Bottom: Overlapping searchlight voxels.

We additionally conducted two additional analyses to examine the robustness of these findings. First, we conducted a searchlight analysis (*25*), in which we trained a PCR model using the same method as we did in the ROI-level modeling for each searchlight (for details, see **Methods**). With LOSO-CV, we identified a set of voxels that showed significant predictions for both pleasure and pain. Their spatial locations largely overlapped with the 7 ROIs that were predictive of both pain and pleasure (Fig. 2B). Second, we tested different PC numbers (or different explained variance) for the PCR modeling, and the results showed similar patterns across PC numbers (**Figs. S3** and **S4**; see **Methods**). Overall, these results showed that our findings were robust across different modeling choices and methods.

### Parsing the affective valence and intensity within the overlapping brain regions

To address the second research question (“Can we identify predictive models of affective valence and intensity within the overlapping brain regions?”), we developed predictive models of intensity and valence, which we hypothesized as two fundamental affective dimensions shared across pain and pleasure (Fig. 3A). We defined the absolute value of pleasantness or unpleasantness rating as an intensity score and the raw signed value of pleasantness (i.e., positive) or unpleasantness (i.e., negative) rating as a valence score. We developed PCR models to predict the intensity or valence scores using the combined data of the capsaicin, chocolate, and control conditions. We used all voxels across the 7 overlapping brain regions for training the intensity and valence models, because a multiple region-based model can yield better prediction performance than single region-based models by capturing information distributed across multiple brain systems (*26*) (for completeness, we also provided region-level intensity and valence modeling results in **Tables S1** and **S2**). Since the overall unpleasantness ratings induced by capsaicin were higher than the overall pleasantness ratings induced by chocolate, we matched the distribution of unpleasantness and pleasantness ratings before prediction by subsampling data and transforming the scores into ranks (±1∼10; please see **Methods**). We tested the resulting models on the independent test dataset (**Study 2**) to examine their generalizability.

**Fig. 3.**
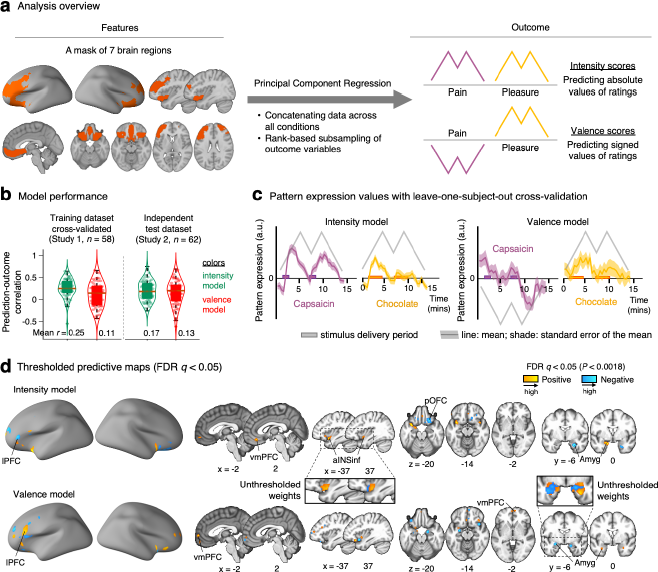
Prediction modeling of affective intensity and valence. **(A)** Overview of the predictive modeling analysis. For the predictive modeling, we used data from the 7 brain regions identified in the previous analysis. We applied principal component regression (PCR) on the concatenated data across all conditions. Before the data concatenation, we conducted the rank-based subsampling based on the outcome variables to minimize the potential bias due to the greater rating magnitude for the capsaicin condition compared to the chocolate condition (for details, see **Methods**). The outcome variable was either affective intensity, which was defined as the absolute values of pleasantness-unpleasantness ratings, or affective valence, which was defined as the raw signed values of the ratings. **(B)** Model performance. The violin plots show the distribution of within-individual prediction-outcome correlations in the training dataset (with LOSO-CV) and the independent test dataset. Colors indicate different models—green for the intensity model and red for the valence model. **(C)** The plots show the time-course of pattern expression values of each model, which were calculated using LOSO-CV. The solid line represents the group average, and the shading represents the standard error of the mean. The different colors indicate the different conditions— purple for capsaicin and yellow for chocolate. The boxes on the time axes indicate the fluid delivery period. **(D)** Predictive weight maps thresholded with FDR *q* < 0.05 based on bootstrap tests (10,000 iterations). Insets show the unthresholded weights for ROIs.

The results showed that both intensity and valence models showed significant prediction performance (Fig. 3B) in the training and independent test datasets. The cross-validated model performance in the training dataset was as the following: for the intensity model, the mean within-individual correlation between actual and predicted ratings was *r* = 0.25, *P* = 2.22 × 10^−16^, bootstrap test, two-tailed, mean mse = 13.061, and for the valence model, *r* = 0.11, *P* = 0.0017, mean mse = 23.860. The models were also predictive of each condition separately (intensity: for the ‘capsaicin’ condition, mean *r* = 0.31, *P* = 4.64 × 10^−8^, mean mse = 19.958, for the ‘chocolate’ condition, mean *r* = 0.23, *P* = 3.53 × 10^−8^, mean mse = 14.859; valence model: for the ‘capsaicin’ condition, mean *r* = 0.12, *P* = 0.0235, mean mse = 35.662, for the ‘chocolate’ condition, mean *r* = 0.08, *P* = 0.0322, mean mse = 27.682), suggesting that these models are sensitive to both conditions. When we tested the models in the independent test dataset (**Study 2**), the intensity and valence models also showed significant prediction: for the intensity model, the mean *r* = 0.17, *P* = 3.79 × 10^−10^, mean mse = 0.0598; for the valence model, the mean *r* = 0.13, *P* = 2.03 × 10^−4^, mean mse = 0.0638. Both models’ pattern expression values showed fluctuations in the same direction with the hypothesized trajectories in the training (Fig. 3C) and independent test datasets (**Fig. S5**).

Although we used ranks instead of original ratings to reduce any potential biases in the models due to different ranges of ratings for the pain and pleasure conditions, we did additional training with the original ratings to examine whether rank-based prediction influenced the model performance. The results showed comparable model performance, mean *r* = 0.27, *P* = 2.22 × 10^−16^, mean mse = 0.0395 for the intensity model, and mean *r* = 0.11, *P* = 7.62 × 10^−4^, mean mse = 0.0651 for the valence model (**Fig. S6A**), suggesting that the rank-based prediction did not have a substantial impact on the model performance. Lastly, we also examined whether using all 48 ROIs for the prediction could have a meaningful effect on the model performance. The models with 48 ROIs also showed comparable model performance compared to the 7 ROI-based models, mean *r* = 0.25, *P* < 2.22 × 10^−16^, mean mse = 13.225 for the intensity model, and mean *r* = 0.09, *P* = 0.0232, mean mse = 24.522 for the valence model (**Fig. S6B**), suggesting that the 7 brain regions are sufficient to predict the affective intensity and valence.

Next, we compared the important voxels between the intensity and valence models based on the thresholded results at FDR *q* < 0.05, two-tailed, from bootstrap tests with 10,000 iterations (for the unthresholded maps, see **Fig. S7**). The intensity and valence maps showed largely non-overlapping spatial patterns of predictive weights (Figs. 3D and 4), suggesting that distinct sub-populations of voxels within the 7 brain regions are important for decoding affective valence versus intensity information. For example, there was a cluster of intensity-predictive voxels in the bilateral ventral anterior insula (Fig. 4A), which has been known as the primary taste cortex (*31*) and encoding the salience of visceral sensation (*31–34*), but there was also a group of valence-predictive voxels in the right ventral anterior insula, mostly non-overlapping with the intensity-predictive voxels (Fig. 4A). Also in the amygdala, the intensity-predictive voxels were mainly located in the ventral part of the right basolateral amygdala with negative weights, and in the dorsal part of the left basolateral amygdala with positive weights (Figs. 3D and 4B). In contrast, the valence-predictive voxels were mainly located in the left centromedial and the right superficial amygdala with negative weights. These results are consistent with previous studies that the basolateral amygdala contains a population of neurons encoding valence-general salience (*15, 20*), whereas the centromedial and superficial amygdala regions are important for processing negative valence (*35–37*). The posterior OFC contained intensity-predictive voxels mostly in the right medial part, while valence-predictive voxels in the lateral part (Fig. 4C). This is also supported by the previous finding that the medial part of this region, as a secondary gustatory cortex (*31*), encode magnitude information of value (*22*). For the ventromedial PFC, the intensity-predictive voxels were located in the ventral part, whereas the valence-predictive voxels were located in the dorsal part (Fig. 4D), and for the dorsolateral PFC, the predictive weights of the two models were also largely non-overlapping (Fig. 4E). In sum, these results suggest that the affective intensity and valence are represented by distinct sub-populations of voxels within the 7 brain regions. However, the overall distribution of intensity-versus valence-predictive voxels across brain regions suggests that the insula is preferentially predictive of intensity, while the ventromedial PFC is preferentially predictive of valence (Fig. 4F). To see the relative importance of all voxels for the intensity versus valence models, see **Fig. S7C**.

**Fig. 4.**
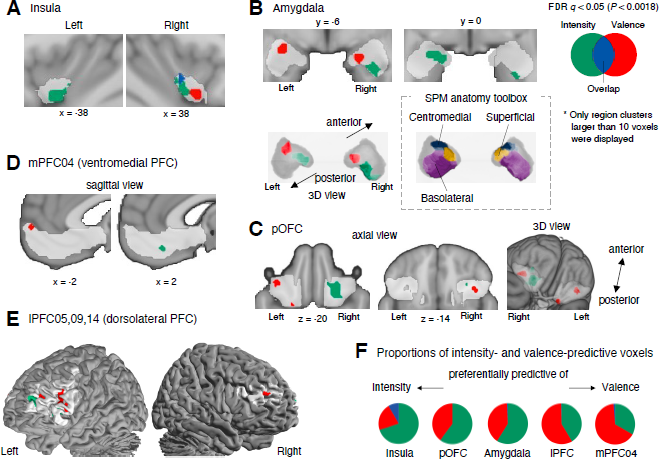
Significant voxels of the intensity and valence models. **(A-E)** To examine whether the significant voxels of the two models are overlapping or not, we examined the exact locations of the significant voxels (FDR *q* < 0.05) by visualizing them with different colors—green for the intensity model, red for the valence model, and blue for the overlapping voxels. Each section shows each ROI: **(A)** insula, **(B)** amygdala, **(C)** posterior OFC (pOFC), **(D)** mPFC04 (ventromedial PFC), and **(E)** lPFC05, 09, and 14 (dorsolateral PFC). We displayed only the clusters that include more than 10 significant voxels. **(F)** The pie charts show the proportions of significant voxels for the intensity model only (green), the valence model only (red), or both models (blue).

### Distinct functional brain networks for the affective intensity and valence models

To situate the locally distinct brain representations of affective intensity and valence in the context of the global functional brain networks, we addressed the last research question (“Which large-scale brain networks are correlated with these predictive maps?”). To this end, we extracted the pattern expression values of intensity and valence models from the ‘control’ condition data using the cross-validated predictive weights. Using these pattern expression values as seeds, we obtained a whole-brain functional connectivity map for each individual. We removed the seed regions (i.e., the mask of 7 brain regions; Fig. 3A) from the resulting maps to help interpretation. Then, we visualized the voxels with significant positive connectivity across participants (FDR *q* < 0.05, one-sample *t*-test, one-tailed) in Fig. 5A. Here we focused only on positive correlations because the negative correlations were difficult to interpret, but for completeness, we also provide the maps with both positive and negative correlations in **Fig. S8**.

**Fig. 5.**
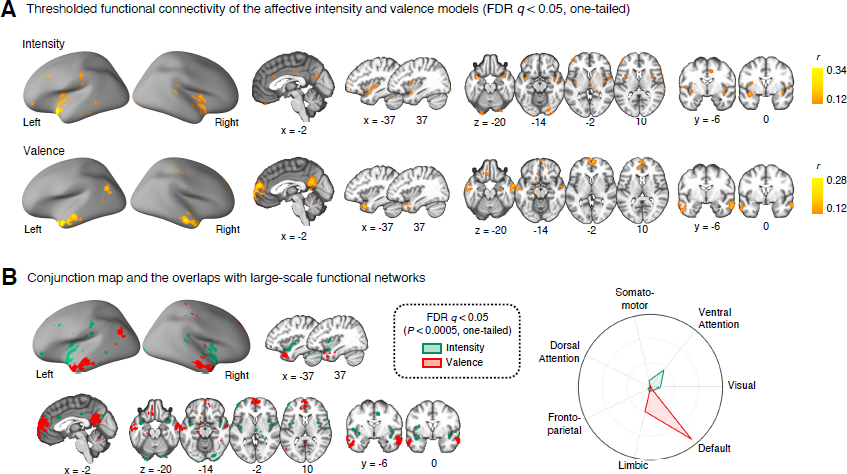
Functional connectivity maps for the intensity and valence models. **(A)** Thresholded functional connectivity patterns using the pattern expression values of intensity and valence models as seeds (FDR *q* < 0.05, one-sample *t*-test, one-tailed). The functional connectivity was calculated using the ‘control’ condition data with LOSO-CV. Here we tested the unidirectional hypothesis (i.e., focusing only on positive correlations) because the negative correlations were difficult to interpret. In addition, we excluded the brain coverage of the predictive models to focus on their relationship with other brain regions. **(B)** Left: Conjunction maps with different colors indicating results of the different models—green for the intensity model and red for the valence model. Right: A radar plot showing the relative proportions of the significant voxels within each large-scale functional network given the total number of voxels within each network.

The results showed that the predictive maps of intensity and valence were correlated with distinct large-scale brain networks. As shown in Fig. 5A, the intensity model showed significant correlations in the insula and the anterior mid-cingulate cortex, the key regions of the ventral attention (or salience) network (*33, 34, 38*). The valence model showed significant correlations in the ventromedial PFC and the posterior cingulate cortex, the key regions of the default mode network. In addition, there were also multiple significant voxels in the OFC and the superior temporal gyrus, which are a part of the limbic network (*23*).

## DISCUSSION

The current study investigated the brain representations of affective valence and intensity in sustained pleasure and pain. We first identified brain regions that contained information about pleasure and pain. The brain regions included the ventromedial and lateral PFCs, the posterior OFC, the ventral anterior insula, and the amygdala. We then developed predictive models of affective intensity and valence with these brain regions, which demonstrated significant prediction performance in both training and independent test datasets. The intensity and valence models were dissociable in their fine-grained spatial patterns of thresholded weights and were associated with the two separate large-scale functional brain networks—the ventral attention network for the intensity information versus the limbic and default mode networks for the valence information. Overall, these results provided neuroimaging evidence of how the brain encodes pleasure and pain based on the general affective dimensions of valence and intensity.

First, our findings support the existence of overlapping brain regions that contain information for both pleasure and pain. Although the relationship between pleasure and pain has been a topic of interest for a long time (*1, 9, 39*), it still remains unclear in which brain regions pleasure and pain are integrated and connected. Previous studies have identified some brain regions commonly activated by pleasure and pain (*9*), but most of the studies examined pleasure and pain separately and did not directly compare their brain representations. Our study aimed to fill this gap by inducing both sustained pleasure and pain in each individual and identifying brain regions predictive of ratings for both conditions. Using a multivariate pattern-based information mapping approach, we identified 7 brain regions for pleasure and pain, including the ventromedial and lateral PFCs, posterior OFC, ventral anterior insula, and amygdala. These regions were among the brain regions that have been implicated in the pleasure-pain interaction (*40–42*) and the affective workspace in the brain (*7*).

Second, our results suggest that the two core dimensions of affective experience— affective intensity and valence—are encoded in these overlapping brain regions. We were able to develop predictive models of affective intensity and valence based on the multivariate patterns of the brain regions’ activity, and the predictive models were generalizable across training and independent test datasets. These results are in line with the previous meta-analysis that there exist both valence-specific and valence-general brain regions (*18*). To further validate our findings, it would be interesting to explore whether the predictive models we developed are responsive to pharmacological or contextual manipulation (*41, 43-47*). Such investigations could have significant implications for clinical applications and represent a promising direction for future research.

Third, through detailed analyses of the model weights, we found that the affective intensity and valence were represented in spatially distinct areas even within the same anatomical brain regions. In addition, the two predictive models were correlated with distinct large-scale functional brain networks—i.e., the ventral attention network was correlated with the affective intensity model, whereas the limbic and default mode networks were correlated with the valence model. These findings are in line with previous studies suggesting that the ventral attention network is important for detecting and identifying important and relevant stimuli given one’s current contexts (*31, 34, 48*), while the limbic and default mode networks are important for modality-general value information (*12, 49*) and subjective affective values (*50–52*). Thus, our results suggest that the affective intensity and valence are processed with distinct brain circuits that are co-localized but connected to distinct large-scale brain systems. This can serve as a guiding hypothesis for more invasive animal studies targeting detailed neural pathways.

Lastly, we also identified some unique brain regions for pleasure and pain. The pain-unique brain regions included the anterior midcingulate cortex (mPFC07), which is known to be preferentially associated with pain among many relevant functions such as negative emotion and cognitive control (*53*). Some lateral PFC regions (lPFC03 and 13) also appeared to be unique to pain. These regions are part of the frontoparietal network, which is known to be important for pain, particularly chronic and sustained pain (*54–56*). The pleasure-unique brain regions included the insular and primary gustatory cortex regions, which could reflect that the affective experience was induced by gustatory stimuli (*57*). It is important to note that the insula anterior long gyrus, which was one of the pleasure-unique insular regions in our results, included the dorsal posterior part of the insular cortex, but this region is well-known for its involvement in pain processing (*58*). However, in our univariate voxel-wise analysis using TR-level data, the dorsal posterior insula appeared to track within-individual changes in ratings for pain (**Fig. S9**), while the region that tracked ratings for pleasure was located at a more anterior part of the insula.

This study also has some limitations. First, although our information mapping and predictive modeling analyses showed significant decoding performances, their effect sizes were small to medium. Though this is not ideal, we reasoned that the small to medium effect sizes might be acceptable in our cases. For example, the main goal of region-level information mapping was to identify the brain regions that contain information about pleasure and pain, not to develop predictive models with the highest effect sizes. In addition, the predictive models of affective intensity and valence showed significant prediction performance even in the independent test dataset. Thus, our results demonstrated that the predictive models were robust despite their small to medium effect sizes. Second, we only used the brain activation patterns for our predictive modeling. Since the experience of sustained pleasure and pain may induce global changes that persist for an extended period of time, an alternative approach could be to utilize patterns of functional connectivity, as we did in our previous study (*56*). Though we focused on the activation patterns to identify brain regions for pleasure and pain in the current study, we could use both activation and connectivity patterns to maximize the prediction performance. This possibility should be examined in future studies. Finally, this study only used taste stimuli. Future studies should test diverse modalities to induce pleasure and pain so that they can investigate more modality-general representations of pleasure and pain.

Overall, the current study provides new insights into the brain representations of affective valence and intensity across pleasure and pain, promoting the system-level understanding of human affective experiences.

## AUTHOR CONTRIBUTIONS

S.A.L. and C.-W.W. conceived, designed, and conducted the experiment, and J.H. and M.C. developed the computer-controlled MR-compatible gustometer for the experiment. S.A.L., J.-J.L., and C.-W.W. analyzed the data, interpreted the results, and wrote the manuscript. J.-J.L., T.D.W. and C.-W.W. edited the manuscript and provided supervision.

## ACKNOWLEDGMENTS

This work was supported by IBS-R015-D1 (Institute for Basic Science; to C.-W.W.), 2019R1C1C1004512, 2021M3E5D2A01022515 (National Research Foundation of Korea; C.-W.W.), and 2E30410-20-085 (KIST Institutional Program; C.-W.W.).

## DATA AVAILABILITY

The data used to generate figures including the predictive models will be shared upon publication.

## CODE AVAILABILITY

The codes for generating the main figures will be shared upon publication. In-house Matlab codes for fMRI data analyses are available at https://github.com/canlab/CanlabCore and https://github.com/cocoanlab/cocoanCORE.

## MATERIALS AND METHODS

### Participants

In Study 1, fifty-eight healthy, right-handed participants were included (27 females, M_age_ = 22.81, SD_age_ = 2.83). We recruited participants from the Suwon area in the Republic of Korea. All participants provided written informed consent in compliance with the approval from the institutional review board (IRB) of Sungkyunkwan University. The preliminary eligibility of participants was determined through an online questionnaire, and those who had psychiatric, neurological, or systemic disorders and MRI contraindications were excluded. In Study 2, sixty-two healthy, right-handed participants were included (29 females, M_age_ = 22.69, SD_age_ = 2.66) based on the same recruitment criteria. Study 1 served as a ‘training dataset’ and was used for developing and evaluating predictive models. Study 2 was an ‘independent test dataset’ used only for the evaluation of the predictive models of intensity and valence.

### Experimental materials and design

The fMRI experiment in Studies 1 and 2 consisted of three conditions; (1) ‘control,’ (2) ‘capsaicin’ (i.e., pain), and (3) ‘chocolate’ (i.e., pleasure) conditions. The orders of these three conditions were counterbalanced across participants. To deliver liquids to participants, we used the custom-built 8-channel fluid delivery system (Octaflow II, ALA Scientific Instruments Inc., Westbury, NY) and a custom-built MR-compatible mouthpiece, where plastic tubes were inserted and connected to Tygon tubes to deliver and extract fluids. In addition, we used a suction device (Join Medical, Inc.) to remove fluids from a participant’s oral cavity during the experiment. The use of this suction procedure helped to minimize head movements caused by the swallowing of fluids.

For the ‘control’ condition, we delivered only water at a very slow speed (about 1.25 ml/min) while participants were relaxed and instructed to continuously rate their current state of pleasantness and unpleasantness using an MR-compatible trackball device. For the ‘capsaicin’ and ‘chocolate’ conditions, capsaicin or chocolate fluids were delivered twice during each scan—first after 90 seconds and then after 7 minutes from the beginning of the scan. Water was delivered at all other times. During scans, we asked participants to rate their subjective level of pleasantness or unpleasantness continuously.

For the capsaicin fluid, we used capsaicin-rich hot sauce (Jinmifood, Inc.), which was diluted with water to a concentration of around 20% (v/v). The capsaicin fluid was delivered for a duration of 90 seconds each, with a delivery volume of less than 1 ml for each delivery. For the chocolate fluid, we used hot chocolate powder (Hershey’s, Inc.), which was dissolved in water to a concentration of around 30% (w/v). The chocolate fluid was delivered for a duration of 3 minutes each, with a delivery volume of around 4.5 ml for each delivery. The duration of fluid delivery was different between capsaicin and chocolate to match the duration of induced pleasure and pain experience similar between the two conditions. To minimize the residual effects of the capsaicin and chocolate fluids, we washed out participants’ oral cavity with water after each condition until they reported no lingering taste. In addition, we conducted the structural scan after the capsaicin condition to allow sufficient time for any remaining painful sensation to be alleviated.

### Rating scale

We used a modified version of the general Labeled Magnitude Scale (gLMS) for the pleasantness-unpleasantness rating. The scale was bidirectional with the center of the scale indicating “Not at all” (0) and the extremes of both sides indicating “Strongest imaginable unpleasantness of any kind” (−1) and “Strongest imaginable pleasantness of any kind” (1). The anchors were “Weak” (0.061), “Moderate” (0.172), “Strong” (0.354), and “Very strong” (0.533) on both sides. Before the fMRI scans, we provided a detailed explanation of how to use the rating scale while showing the anchors, but the anchors were not shown on the screen during the scan. To reduce the potential confounding effects of rating directionality, we randomly switched the labels of two rating extremes (i.e., pleasantness and unpleasantness) across participants.

### fMRI data acquisition

Whole-brain fMRI data were acquired on a 3T Siemens Prisma scanner in the Center for Neuroscience Imaging Research at Sungkyunkwan University. High-resolution T1-weighted structural images were acquired. Functional EPI images were acquired with TR = 460 ms, TE = 27.2 ms, multiband acceleration factor = 8, field of view = 220 mm, 82 × 82 matrix, 2.7 × 2.7 × 2.7 mm^3^ voxels, 56 interleaved slices, number of volumes = 1893. We used Matlab (Mathworks) and Psychtoolbox (http://psychtoolbox.org/) for the stimulus presentation and behavioral data acquisition.

### fMRI data analysis

Structural and functional MRI data were preprocessed using our in-house preprocessing pipeline (https://github.com/cocoanlab/humanfmri_preproc_bids). This pipeline utilizes preprocessing tools in FSL and SPM12. We removed 18 initial volumes of functional EPI images for image intensity stabilization. Then, we co-registered structural T1-weighted images to the functional images and normalized them to MNI, and the functional EPI images were distortion-corrected, motion-corrected (realigned), normalized to MNI using T1 images with the interpolation to 2 × 2 × 2 mm^3^ voxels, and smoothed with a 5-mm FWHM Gaussian kernel. We then used 25-second time bin regressors convolved with the hemodynamic response function for GLM analysis and obtained 34 beta images. In the GLM analysis, we included nuisance covariates including 24 head motion parameters (x, y, z, roll, pitch, and yaw, their mean-centered squares, their derivatives, and squared derivative), linear drift, outlier indicators, and five principal components of white matter signals and those of ventricle signals. We also winsorized the values that were detected as spatiotemporal outliers (intensity values < median minus 5 SDs or > median plus 5 SDs) and applied a low pass filter with a 0.1 Hz threshold. The resulting beta images were concatenated and rescaled across participants and conditions in the SPM style (i.e., L1 normalization). We used these data as input features for the predictive modeling. We also down-sampled the pleasantness-unpleasantness ratings into 34 time-bins and used them as an outcome variable for the predictive modeling. A graphical illustration of making 34-bin data of fMRI and behavioral data is shown in **Fig. S2A**.

### Region-level information mapping

To identify which brain regions were predictive of pain or pleasure, we conducted predictive modeling based on multivariate patterns of brain activity for different brain regions. We first defined 48 *a priori* regions-of-interest (ROIs) covering the brainstem, subcortical regions, and prefrontal regions, based on previous studies (*28–30*) and the Harvard-Oxford brain atlas. For each ROI, we trained predictive models of pain or pleasure using the principal component regression (PCR) algorithm (*59*). The modeling was based on the concatenated fMRI data across all 34 time-bins, conditions (i.e., ‘capsaicin’ and ‘control’ conditions or ‘chocolate’ and ‘control’ conditions), and participants.

Since the choice of the number of principal components (PCs) can significantly affect the prediction performance, we used the same number of PCs for all 48 regions. To determine the number of PCs *a priori*, we first averaged all the time-bin data for each condition and concatenated the averaged fMRI data across the ‘capsaicin’ and ‘control’ conditions or across the ‘chocolate’ and ‘control’ conditions, obtaining two images per participant, resulting in a total of 116 images (= 2 images × 58 participants) for each condition. Then, we applied principal component analysis (PCA) to the concatenated data to find the minimum required number of components to explain > 75% of the total variance. We repeated this procedure for each ROI, obtained a total of 96 different PC numbers (= 48 ROIs × 2 conditions), and selected the median of the PC numbers, which was 13 PCs (graphical illustration is provided in **Fig. S2, B** and **C**). We also tested different PC numbers for different levels of explained variance (i.e., 65%, 70%, 75%, 80%, and 85%), which showed similar performances with gradual changes (**Figs. S3** and **S4**).

We evaluated the mean within-individual correlation between predicted and actual outcomes (i.e., pleasantness-unpleasantness ratings) with the leave-one-subject-out cross-validation (LOSO-CV) and examined their statistical significance using bootstrap tests with 10,000 iterations. The ROIs that showed statistically significant predictions for both pleasantness and unpleasantness ratings (FDR-corrected) were selected for further analyses of affective intensity and valence.

### Searchlight analysis

To examine the robustness of the region-level information mapping results, we repeated the predictive modeling with searchlights. For this, we created a searchlight with a radius of five voxels and scanned it throughout the whole brain with a step size of four voxels. The number of PCs was the same with the region-level predictive modeling (= 13) for consistency. We developed a PCR model for each searchlight region, evaluated the prediction performance with LOSO-CV, and added this value to cubes which had one side of four voxels and were located at the center of each searchlight across the whole brain with no overlapping voxels between cubes. We smoothed the performance maps of pain and pleasure models with a 3-mm FWHM Gaussian kernel, and thresholded results based on the bootstrap test with 10,000 iterations and FDR correction.

### Predictive modeling of affective intensity and valence

To examine how the 7 brain regions for pain and pleasure encode two core affective dimensions—affective intensity and valence, we first defined affective intensity as the absolute values of pleasantness-unpleasantness ratings and affective valence as the raw signed values of pleasantness-unpleasantness ratings. Using these as outcomes, we developed the PCR models to predict affective intensity and valence across all conditions (i.e., capsaicin, chocolate, and control) and participants. We used all the voxels within the 7 brain regions for this modeling. However, there was an issue in using the concatenated ratings across different conditions—the rating magnitudes for the capsaicin condition were greater than those for the chocolate condition, creating a skewness of the data distribution and thus introducing a potential bias in the modeling.

To minimize the potential bias, we subsampled the data with a stratified sampling method to match the numbers of positive (i.e., pleasantness) and negative (i.e., unpleasantness) rating data. In more detail, we first split the rating data into ten equidistant bins and transformed them into positive and negative ranks (i.e., 1, 2, …, 10 and −1, −2, …, −10), except for zeros. We then compared the frequency of positive and negative values in a bin-by-bin manner, i.e., the number of data points within the bin +1 vs. −1, +2 vs. −2, …, and matched the frequency by random subsampling of data from the bin that had more data. Consequently, the distributions of the positive and negative values became symmetric. We developed the intensity and valence models using these subsampled data.

The other procedures of predictive modeling were identical to the region-level information mapping. The number of PCs that explained 75% of the total variance was 123 for both intensity and valence models, and thus we chose to use 123 PCs for the PCR modeling. Prediction performances of the intensity and valence models were evaluated using LOSO-CV. To examine the robustness of these two models, we also tried (1) developing PCR models to predict the original rating values instead of the ranks and (2) using the whole 48 ROIs (**Fig. S6**). The procedure of predictive modeling of intensity and valence for each of the 7 overlapping brain regions was also identical, using 31 PCs based on the same method used to choose the number of PCs in the region-level information mapping (**Tables S1** and **S2**).

The model predictions (or model responses) were calculated as the dot product of vectorized brain activity with predictive weights, which we termed pattern expression (i.e., the expression of the multivariate pattern of a predictive model).

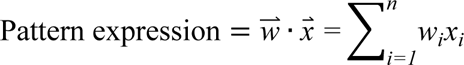

where *n* is the number of voxels, *w* is the predictive weights, and *x* is the fMRI data. To minimize bias in the estimation of model performance, we used LOSO-CV—i.e., pattern expression values of one participant were calculated using the predictive weights trained on the data after excluding the participant’s data. We calculated the prediction-outcome correlation (i.e., a correlation between the pattern expression values and actual outcomes) for each individual and conducted bootstrap tests (i.e., resampling with replacement) on the correlation values with 10,000 iterations. To identify the voxels that reliably contributed to prediction, we conducted bootstrap tests with 10,000 iterations with using the same PC number (= 123). To test the generalizability of the models, we tested them on an independent test dataset (Study 2) with no *ad hoc* tuning of the predictive weights. We calculated the prediction-outcome correlation for each participant and conducted bootstrap tests.

### Whole-brain functional connectivity analysis of the intensity and valence models

To examine which brain regions and networks are correlated with the predictive models of affective intensity and valence, we calculated the correlations between the pattern expression values and the whole-brain fMRI data. We used the data of the control condition of Study 1 for this analysis because it involved no painful or pleasant stimulation and thus reflected the intrinsic brain activity. For each participant, this analysis yielded the whole-brain functional connectivity maps with pattern expression values as seeds (*60*). We then conducted *t*-test to test whether the functional connectivity was significantly greater than zero across participants (i.e., positive connectivity) with FDR correction. We did not consider negative connectivity because anti-correlations were difficult to interpret. However, for completeness, we also obtained the thresholded functional connectivity maps with both positive and negative correlations in **Fig. S8**. We also calculated how many suprathreshold voxels were included in each large-scale functional brain network (*61*).

