## Supplementary Information for "Brain Representations of Affective Valence and Intensity in Sustained Pleasure and Pain"

**This PDF file includes:**

1. Figs. S1 to S9
2. Table S1 to S2
3. References

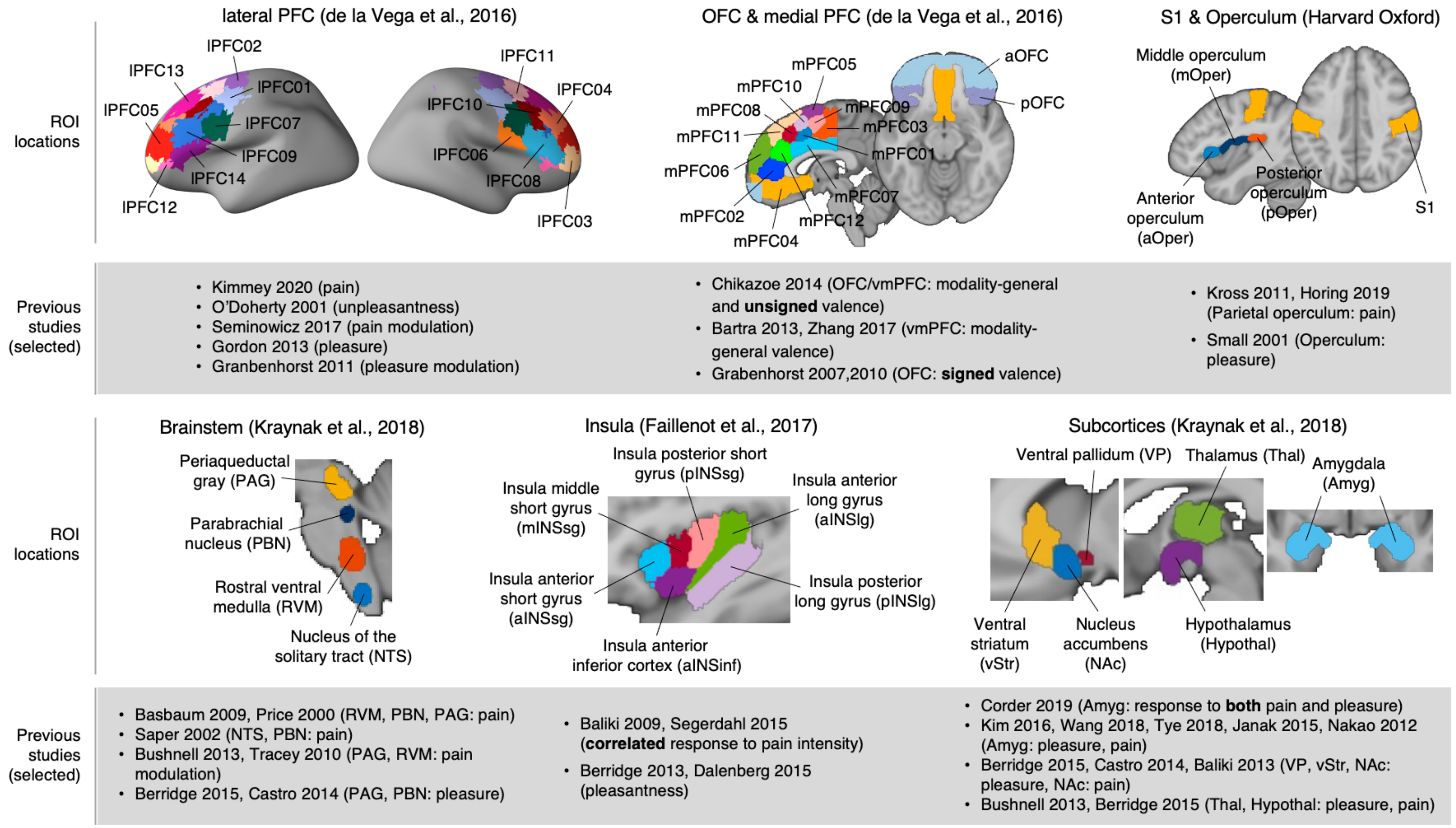


**Fig. S1. 48 *a priori* regions-of-interest (ROIs) (related to Figs. 1-2).**
We selected the 48 brain regions *a priori* that have been reported to be responsive to either pain or pleasure. First, the lateral prefrontal cortex has been known to be related to pain, pleasure, and their modulation(*1-5*). Second, the orbitofrontal and medial prefrontal cortices have been heavily studied in terms of their modality-general valence processing(*6-10*). Third, the primary somatosensory cortex (S1) and operculum have been reported as pain and pleasure-responsive regions(*11-13*). Fourth, the brainstem regions including the nucleus of the solitary tract, rostral ventral medulla, parabrachial nucleus and periaqueductal gray, are known to be part of ascending and descending pain pathways as well as of pleasure pathways(*14-20*). Fifth, the insular cortex has been known to encode pleasantness(*21, 22*), and particularly, the dorsal posterior insula (especially near the parietal operculum) has shown to be correlated with pain intensity(*23, 24*). Sixth, the ventral striatum, nucleus accumbens, ventral pallium, thalamus and hypothalamus are also

known to be important for processing pain and pleasure(*17, 19, 20, 25*). Lastly, the amygdala has been heavily studied due to its response both to pain and pleasure(*26-31*). We used the defined ROIs from previous studies, for brainstem and subcortical regions, ref. (*32*), for insular regions, ref. (*33*), and for lateral and medial prefrontal cortices (PFC), ref. (*34*). We used the Harvard-Oxford brain atlas for S1 and opercular regions.

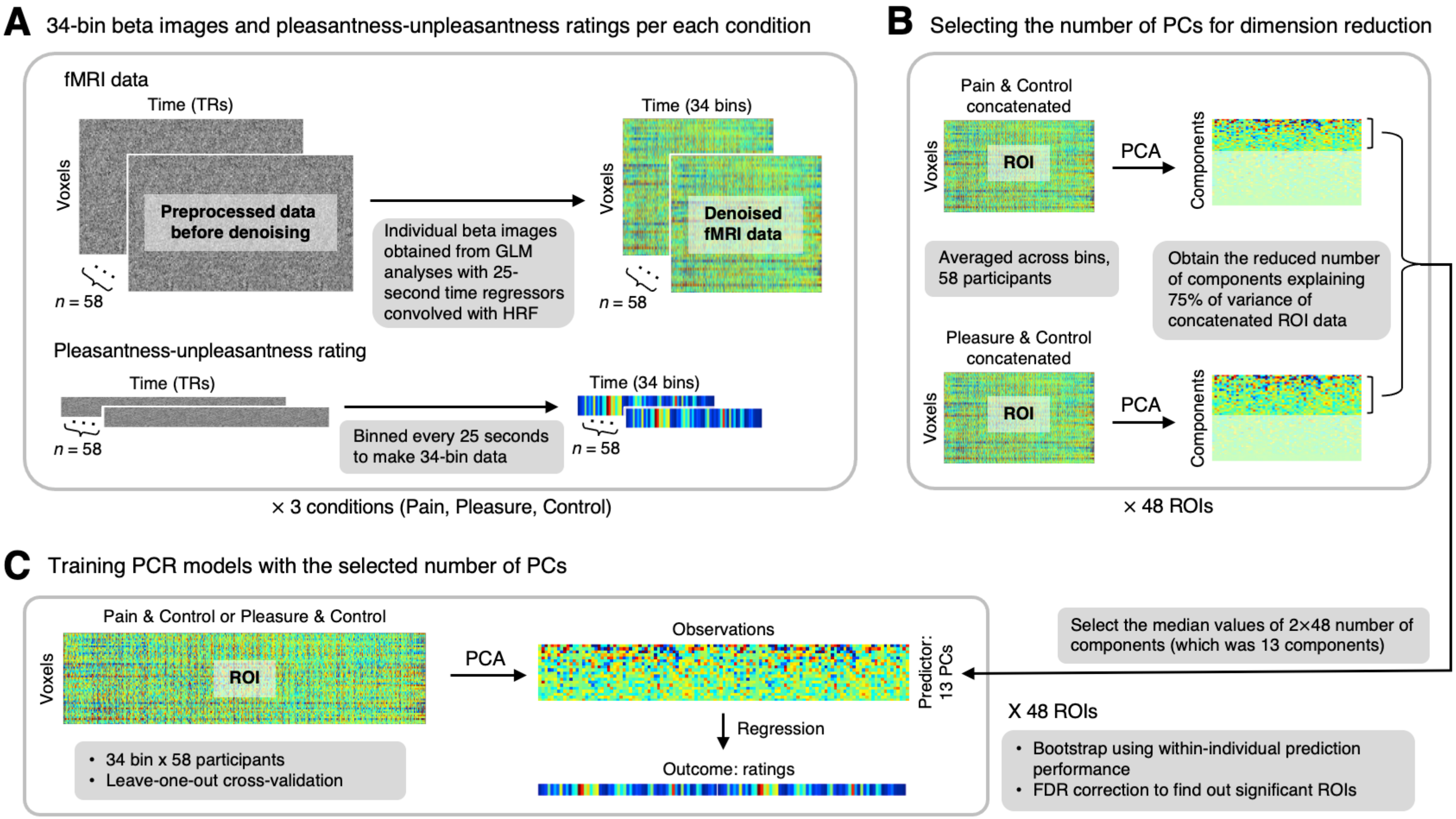


**Fig. S2. Analysis overview of region-level predictive modeling (related to Fig. 2).**
**(A)** For the predictive modeling, we used 34-bin beta images of fMRI data as inputs. To obtain 34-bin beta images, we conducted general linear modeling (GLM) analyses with 25-second time-bin regressors convolved with the hemodynamic response function. We also included nuisance covariates (24 head motion parameters, linear drift, outlier indicators, and five principal components of white matter and ventricle signals) in the GLM analyses. We also averaged the pleasantness-unpleasantness ratings into 34 time-bins. **(B)**

Given that the choice of the number of principal components (PCs) can influence the model training, we aimed to use the same number of PCs for all 48 regions. To determine the number of PCs, we first averaged all the time-bin data for each condition and concatenated the averaged fMRI data across the capsaicin and control conditions or across the chocolate and control conditions, obtaining two images per participant, resulting in a total of 116 images (= 2 images × 58 participants) for each condition. Then, we applied principal component analysis (PCA) to the concatenated data to find the minimum required number of components to explain > 75% of the total variance. We repeated this procedure for each ROI, obtained 96 different PC numbers (= 48 ROIs × 2 conditions), and selected the median of the PC numbers, which was $n = 13$ components. **(C)** Using this number of PCs, we applied the principal component regression (PCR) for each ROI to predict ratings for either the capsaicin and control conditions or the chocolate and control conditions, based on concatenated fMRI data across all the time-bins, conditions, and participants (i.e., 34 bins × 2 runs × 58 participants). We evaluated the mean within-individual correlation between predicted and actual outcomes with the leave-one-subject-out cross-validation and examined their statistical significance using bootstrap tests with 10,000 iterations. We used the false discovery rate (FDR) $q < 0.05$ to correct for the multiple comparisons.

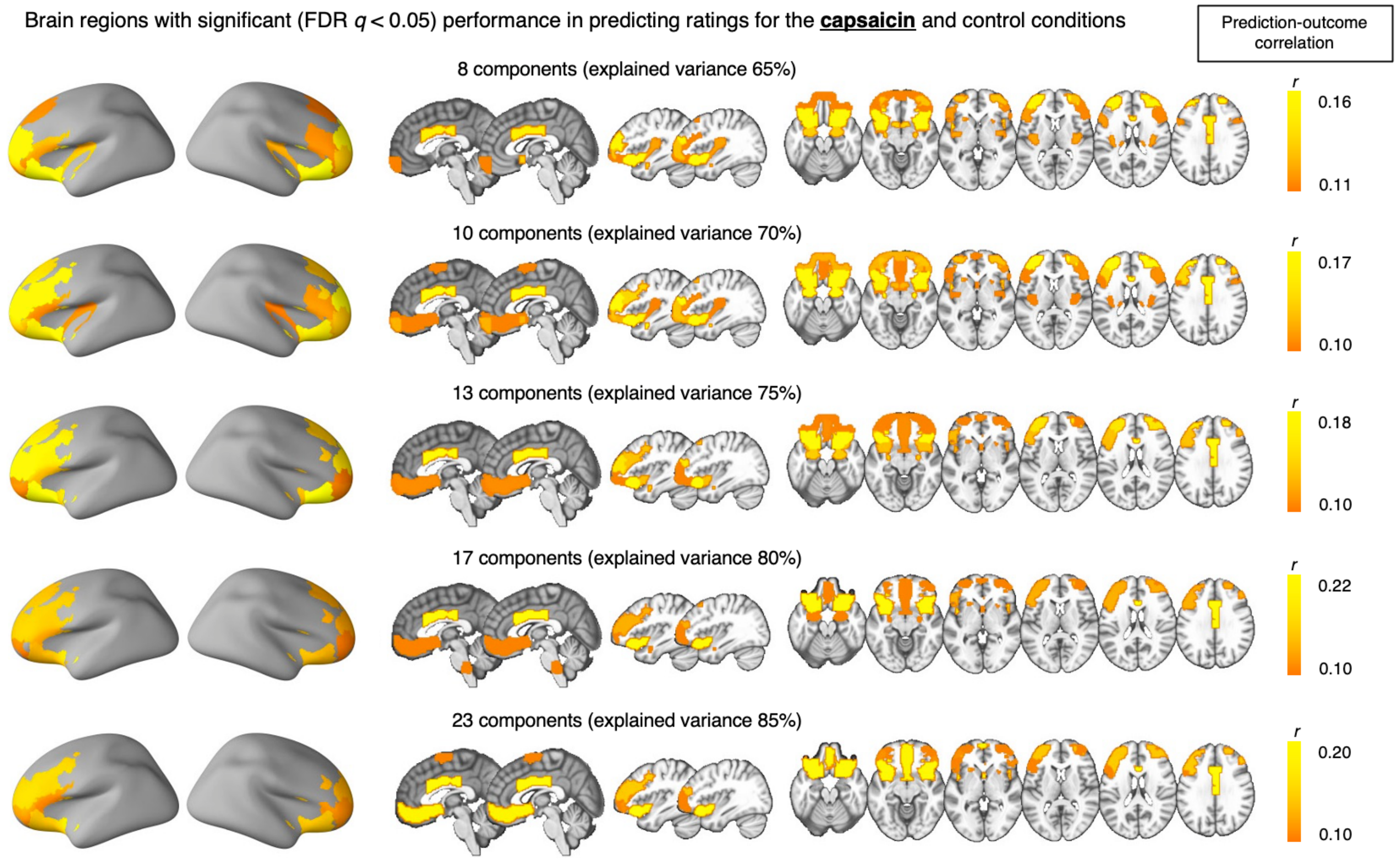


**Fig. S3. Region-level predictive modeling for the capsaicin and control conditions with different numbers of principal components (related to Fig. 2).**

Though we used 13 PCs for main analyses, we also tested different numbers of PCs to examine how robust our results were. We varied the number of PCs (8~23) with different levels of explained variance (i.e., 65%, 70%, 75%, 80%, and 85%) and examined which brain regions showed significant prediction performance in predicting ratings for the capsaicin and control conditions. The brain maps show regions with significant prediction performance thresholded at FDR $q < 0.05$.

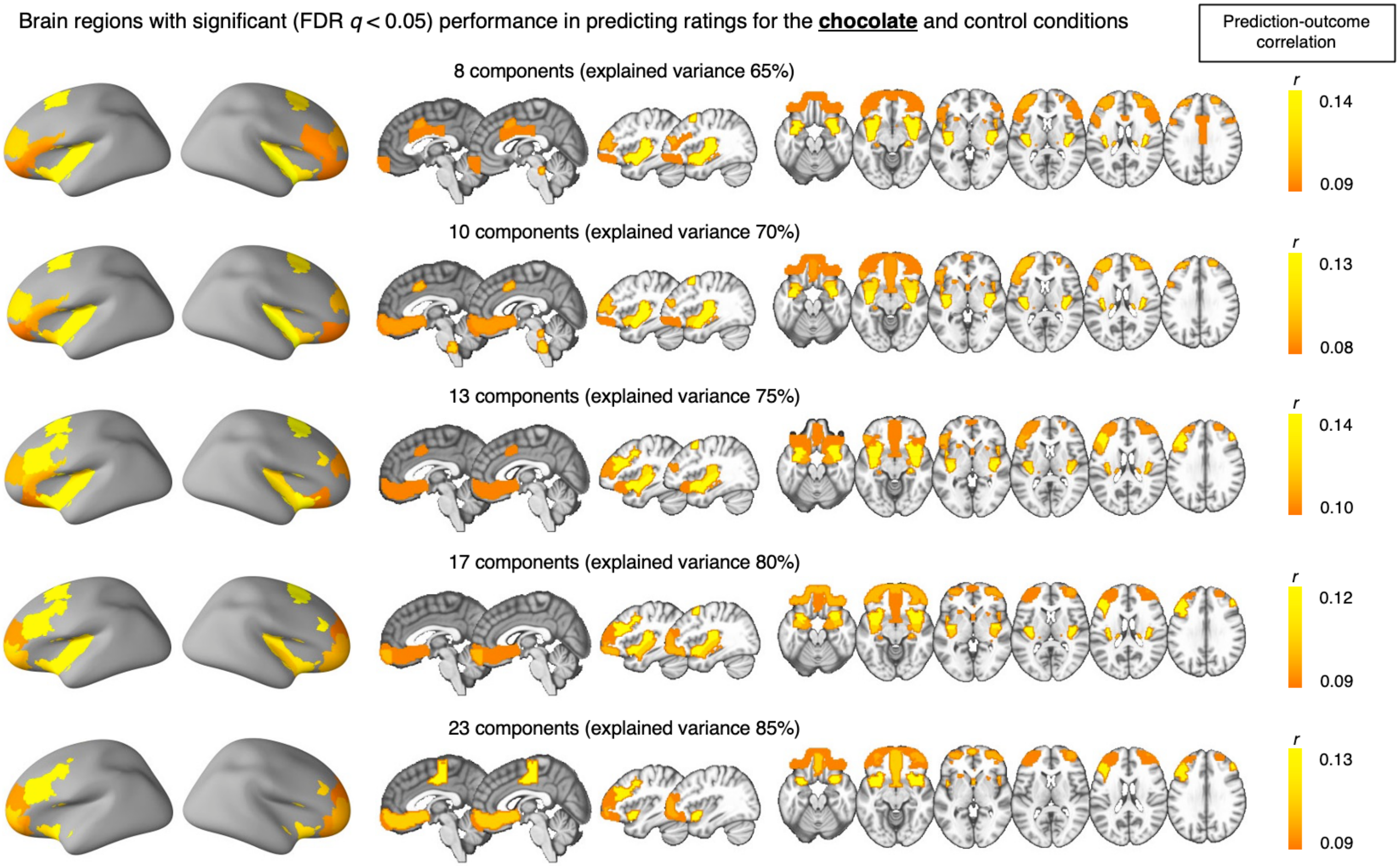


**Fig. S4. Region-level predictive modeling for the chocolate and control conditions with different numbers of principal components (related to Fig. 2).**
Same as Fig. S3, but the results of predicting ratings for the chocolate and condition conditions.

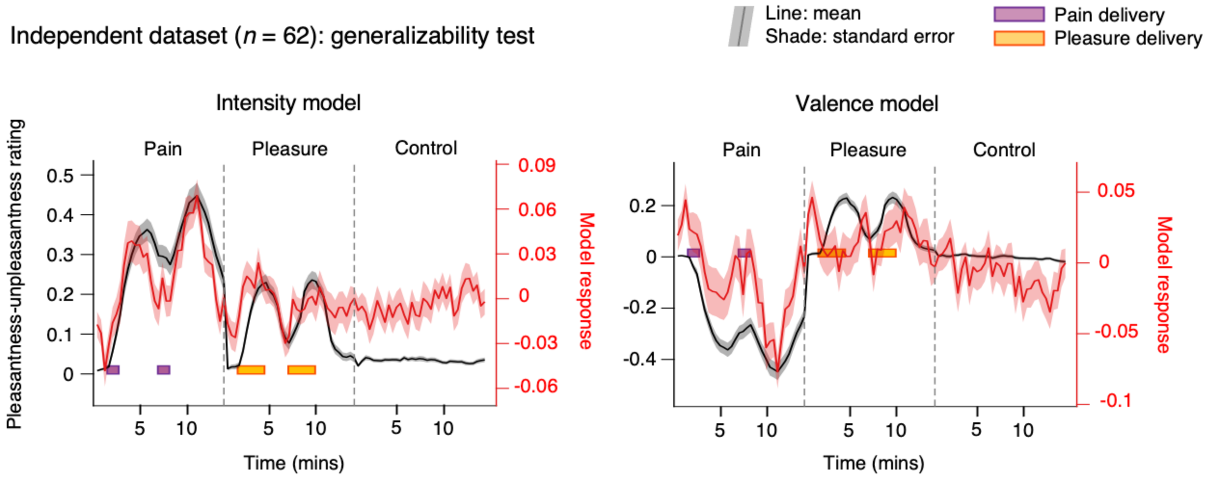


**Fig. S5. Testing intensity and valence models on the independent dataset (Study 2, *n* = 62; related to Fig. 3).**

We tested whether the intensity and valence models also showed the time-course of pattern expression values similar to the training dataset results. The solid line represents the group average, and the shading represents the standard error of the mean. The colored boxes indicate the fluid delivery period (purple for capsaicin and yellow for chocolate).

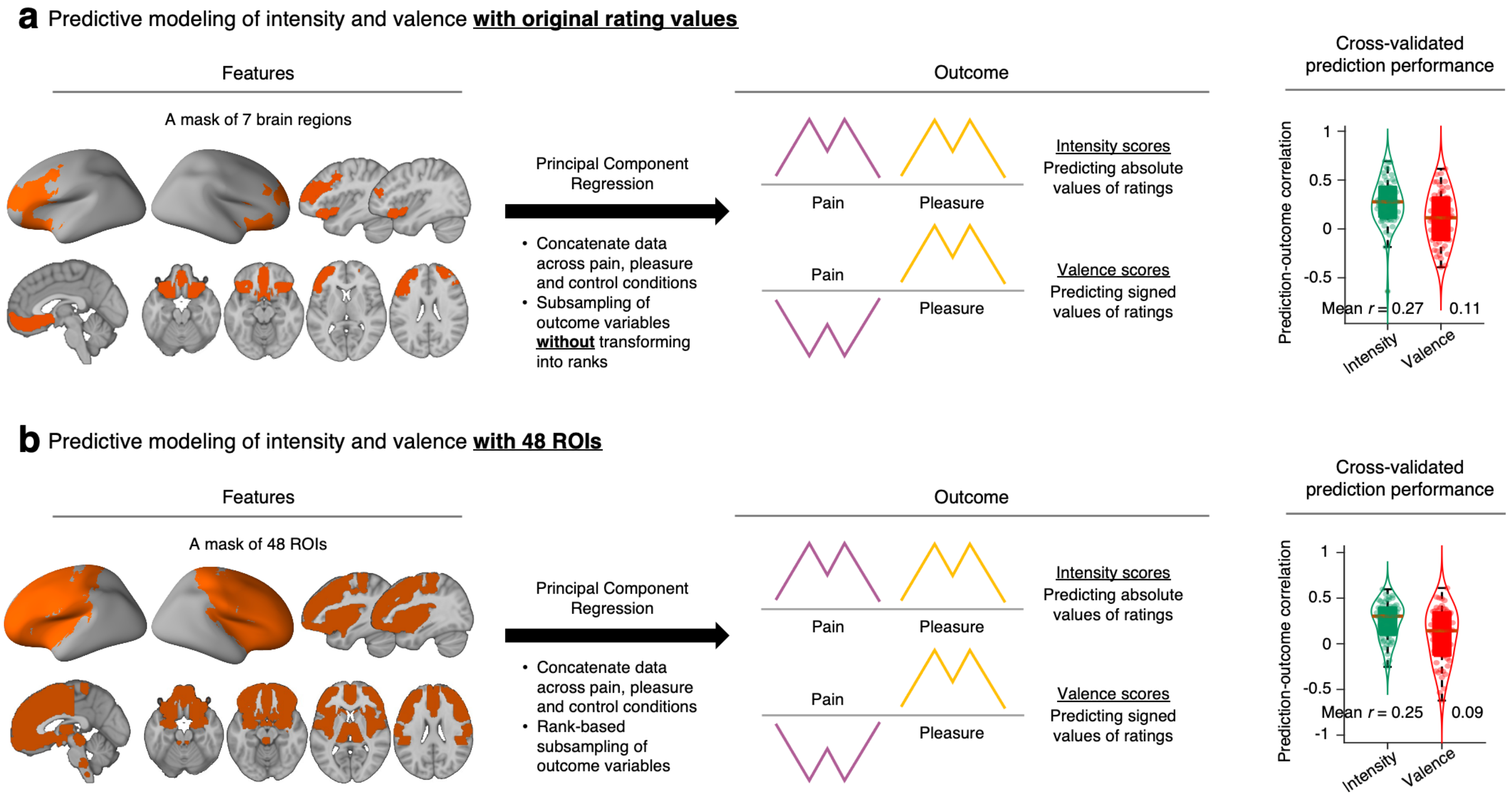


**Fig. S6. Predictive modeling of affective intensity and valence using original rating values and the whole 48 ROIs (related to Fig. 3).**
**(A)** When we used the original rating values as the outcome, instead of the rank values, the mean within-individual prediction-outcome correlation for the intensity model was $r = 0.27$, $P = 2.22 \times 10^{-16}$, bootstrap test, two-tailed, mean of mean squared error (mse) = 0.0395, and for the valence model, $r = 0.11$, $P = 7.62 \times 10^{-4}$, bootstrap test, two-tailed, mean mse = 0.0651. **(B)** When we used 48 ROIs instead of 7 ROIs, the mean within-individual prediction-outcome correlation for the intensity model was $r = 0.25$, $P < 2.22 \times 10^{-16}$, bootstrap test, two-tailed, mean mse = 13.225 and for the valence model, $r = 0.09$, $P = 0.0232$, bootstrap test, two-tailed, mean mse = 24.522.

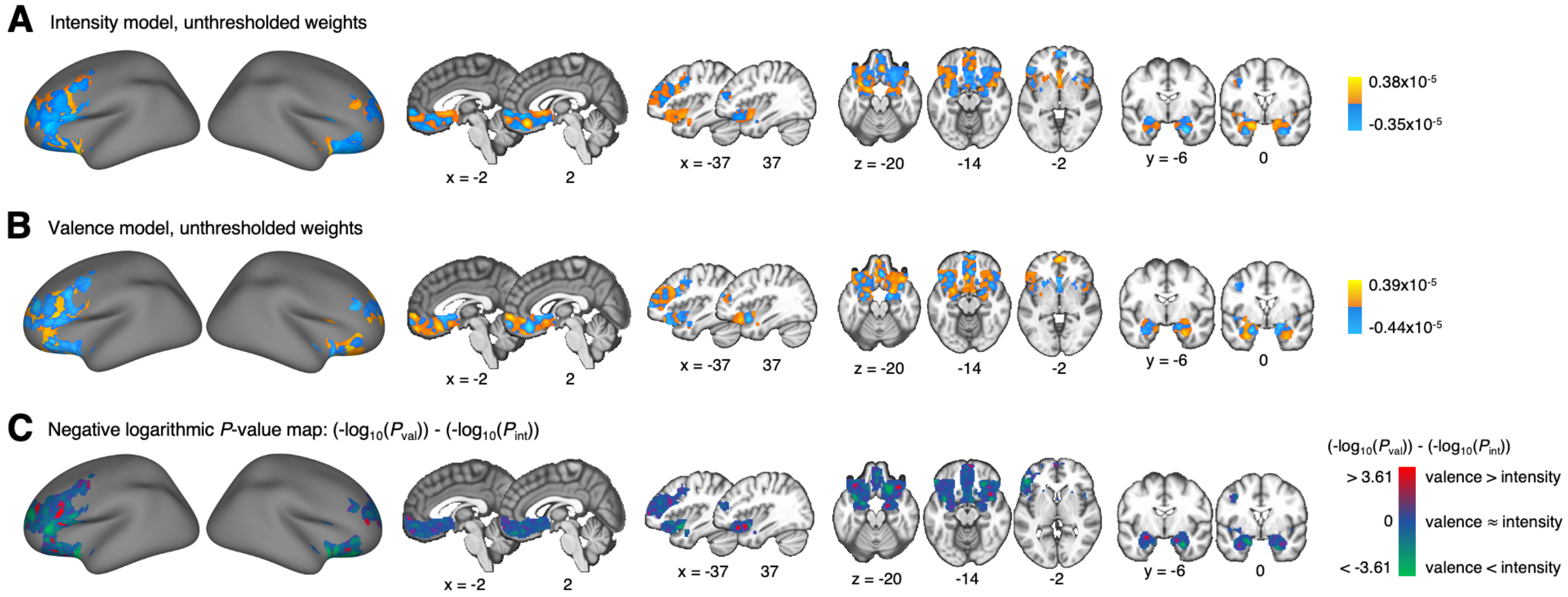


**Fig. S7. Unthresholded predictive weights of intensity and valence models (related to Fig. 3).**
**(A)** Unthresholded predictive weights of the intensity model. **(B)** Unthresholded predictive weights of the valence model. **(C)** The relative importance of the voxels for the intensity versus valence models. Here we defined the importance of the voxels as the negative logarithmic of *p*-values to base 10 from bootstrap tests. The brain regions with lower *p*-values for the intensity model were colored green, and the brain regions with lower *p*-values for the valence model were colored red.

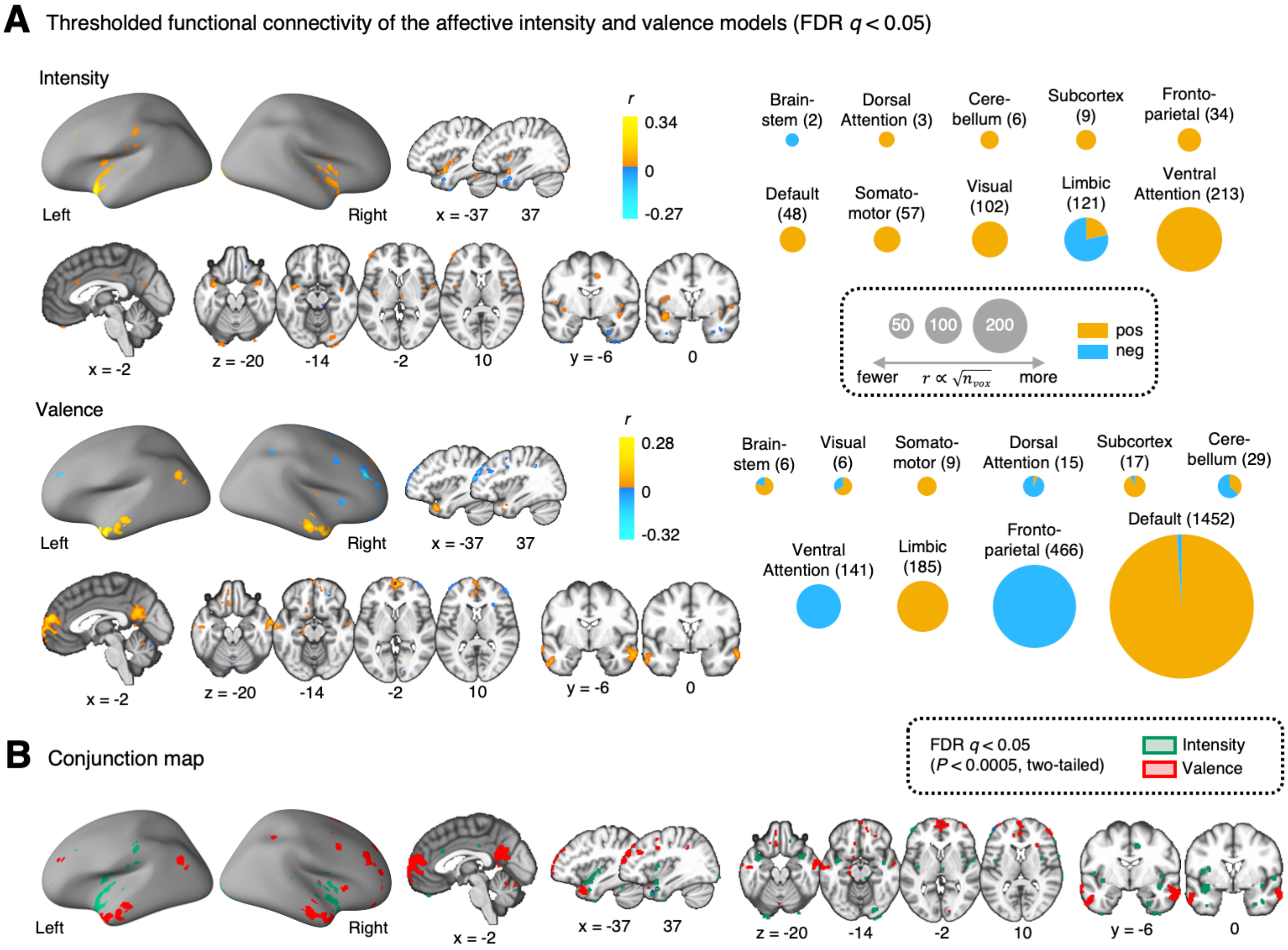


**Fig. S8. Functional connectivity maps for the intensity and valence models including both positive and negative correlations (related to Fig. 5).**
**(A)** Left: Thresholded functional connectivity maps of intensity and valence models including both positive and negative correlation values (FDR $q < 0.05$, one-sample $t$-test, two-tailed). We excluded the brain coverage of the predictive models to focus on their

relationship with other brain regions. Right: The pie charts showed the proportions of voxels that survived after thresholding given each large-scale functional brain network. The radius of each pie chart is proportional to the total number of voxels; the more voxels in each brain network survive, the bigger the pie chart is. The actual numbers of all survived voxels in each brain network are indicated in parentheses. **(B)** A conjunction map of the intensity and valence model-based connectivity maps; green for the intensity model and red for the valence model.

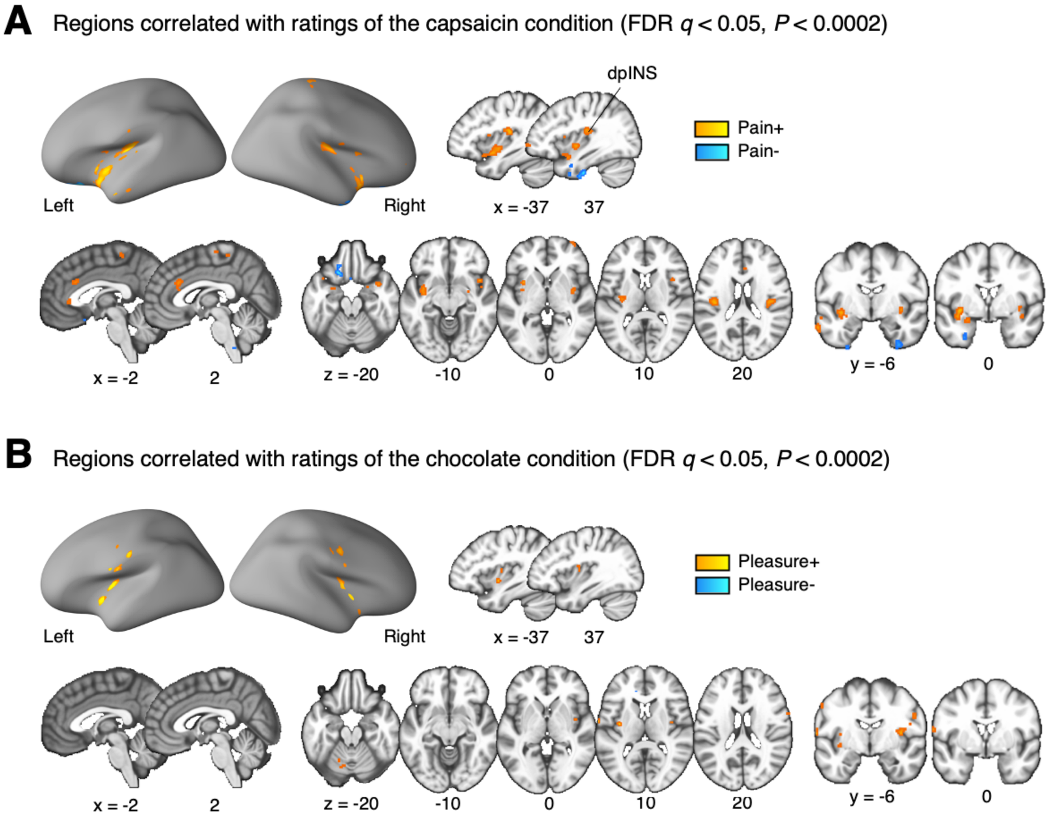


**Fig. S9. Univariate GLM analysis using TR-level fMRI and pleasantness-unpleasantness ratings (related to Fig. 2).**
**(A-B)** To examine which voxels were correlated with the ratings during the capsaicin versus chocolate conditions, we conducted a univariate general linear modeling (GLM) analysis using TR-level fMRI and behavioral data. We used the TR-level pleasantness-unpleasantness ratings convolved with the canonical hemodynamic response function as a regressor for **(A)** the 'capsaicin' and **(B)** the 'chocolate' conditions. We also included the nuisance covariates (24 head motion parameters, linear drift, outlier indicators, and five principal components of white matter and ventricle signals) to remove nuisance effects in each fMRI condition. We then conducted one-sample *t*-tests for the beta coefficients with FDR correction for multiple comparisons.

**Table S1. Predicting affective intensity using individual 7 overlapping brain regions (related to Fig. 3).**

| Brain regions | All conditions | | 'Capsaicin' condition | | 'Chocolate' condition | |
|---|---|---|---|---|---|---|
| | Mean $r$ | $P$ | Mean $r$ | $P$ | Mean $r$ | $P$ |
| Amygdala | 0.15 | $3.02 \times 10^{-5*}$ | 0.20 | $3.91 \times 10^{-6*}$ | 0.18 | $2.95 \times 10^{-6*}$ |
| Insula anterior inferior cortex | 0.18 | $8.16 \times 10^{-6*}$ | 0.23 | $2.81 \times 10^{-6*}$ | 0.19 | $1.44 \times 10^{-5*}$ |
| lPFC05 | 0.11 | $0.0021^{*}$ | 0.17 | $0.0006^{*}$ | 0.03 | 0.4568 |
| lPFC09 | 0.07 | 0.0546 | 0.07 | 0.1243 | -0.001 | 0.9417 |
| lPFC14 | 0.10 | $0.0012^{*}$ | 0.12 | 0.0064 | 0.05 | 0.1742 |
| mPFC04 | 0.10 | 0.0044 | 0.17 | $0.0004^{*}$ | 0.09 | 0.0141 |
| Posterior OFC | 0.15 | $1.27 \times 10^{-5*}$ | 0.22 | $1.93 \times 10^{-5*}$ | 0.15 | $0.0011^{*}$ |

*Note.* The procedure of predictive modeling using each of the 7 brain regions was identical to that of using all 7 brain regions. The number of PCs that explained 75% of the total variance across 7 brain regions was 31, also an identical procedure used in the region-level information mapping (see **Methods**). lPFC: lateral prefrontal cortex, mPFC: medial prefrontal cortex, OFC: orbitofrontal cortex. $^{*}q < 0.05$, false discovery rate corrected, one-sample *t*-test, two-tailed.

**Table S2. Predicting affective valence using individual 7 overlapping brain regions (related to Fig. 3).**

| Brain regions | All conditions | | 'Capsaicin' condition | | 'Chocolate' condition | |
|---|---|---|---|---|---|---|
| | Mean *r* | *P* | Mean *r* | *P* | Mean *r* | *P* |
| Amygdala | 0.04 | 0.2554 | 0.05 | 0.2708 | 0.05 | 0.2162 |
| Insula anterior inferior cortex | 0.07 | 0.1287 | -0.03 | 0.4397 | 0.07 | 0.0931 |
| lPFC05 | 0.12 | 0.0009 | 0.14 | 0.0018 | 0.04 | 0.3352 |
| lPFC09 | 0.12 | 0.0002* | 0.18 | 0.0001* | 0.01 | 0.7520 |
| lPFC14 | 0.10 | 0.0542 | 0.08 | 0.0985 | 0.04 | 0.2676 |
| mPFC04 | 0.06 | 0.0881 | 0.04 | 0.4764 | 0.05 | 0.2082 |
| Posterior OFC | 0.06 | 0.2243 | 0.05 | 0.2555 | 0.04 | 0.2637 |

*Note.* Same as Table S1, but the results of affective valence prediction. lPFC: lateral prefrontal cortex, mPFC: medial prefrontal cortex, OFC: orbitofrontal cortex. $^{*}q < 0.05$, false discovery rate corrected, one-sample $t$-test, two-tailed.
